# Uncovering the Structure and Function of Microbial Communities Formed During Periodic Tilling of TNT and DNT Co-Contaminated Soils

**DOI:** 10.1101/2020.12.12.420737

**Authors:** Saeed Keshani-Langroodi, Yemin Lan, Ben Stenuit, Gail Rosen, Joseph B. Hughes, Lisa Alvarez-Cohen, Christopher M. Sales

## Abstract

1.

Environmental contamination by 2,4,6-trinitrotoluene (TNT), historically the most widely used secondary explosive, is a long-standing problem in former military conflict areas and at manufacturing and decommissioning plants. In field test plots at a former explosives manufacturing site, removal of TNT and dinitrotoluenes (DNTs) was observed following periods of tillage. Since tilling of soils has previously been shown to alter the microbial community, this study was aimed at understanding how the microbial community is altered in soils with historical contamination of nitro explosives from the former Barksdale TNT plant. Samples of untilled pristine soils, untilled TNT-contaminated soils and tilled TNT-contaminated soils were subjected to targeted amplicon sequencing of 16S ribosomal RNA genes in order to compare the structure of their bacterial communities. In addition, metagenomic data generated from the TNT tilled soil was used to understand the potential functions of the bacterial community relevant to nitroaromatic degradation. While the biodiversity dropped and the *Burkholderiales* order became dominant in both tilled and untilled soil regardless of tillage, the bacterial community composition at finer taxonomic levels revealed a greater difference between the two treatments. Functional analysis of metagenome assembled genome (MAG) bins through systematic review of commonly proposed DNT and TNT biotransformation pathways suggested that both aerobic and anaerobic degradation pathways were present. A proposed pathway that considers both aerobic and anaerobic steps in the degradation of TNT in the scenario of the tilled contaminated soils is presented.

**Importance:** In this study, TNT and DNT removal has been observed in field-scale experiments following periodic tilling of historically contaminated soils. The microbial community structures of uncontaminated pristine soils, untilled contaminated soils, and tilled contaminated soils were investigated using high-throughput sequencing platforms. In addition, shotgun metagenome libraries of samples from tilled contaminated soils were generated. The results indicated that a significant shift of the bacterial community at the family level between tilled and untilled contaminated soils, with tilled soils being dominated by *Alcaligenaceae* and untilled soils by *Burkholderiacea.* In-depth metagenomic analysis of samples from tilled contaminated soils, indicate the presence of genes that encode for enzymes that potentially could lead to mineralization of TNT and DNT under mixed aerobic and anaerobic periods.

## 3. Introduction

2,4,6-Trinitrotoluene (TNT), an anthropogenic nitroaromatic explosive synthesized by successive nitration of toluene, is ranked high in the Priority List of Hazardous Substances (ATSDR 2000). Manufacturing plants and military sites are major sources of environmental pollution of explosives (Stenuit and Agathos 2010, Stenuit and Agathos 2011, Stenuit and Agathos 2011, USEPA 2014). In former manufacturing plants, high concentration of TNT in soils (over 100 mg/kg soil) can persist for long periods of time (Hewitt, Jenkins et al. 2003, Pennington, Jenkins et al. 2006). In addition, mononitrotoluenes (MNTs) and dinitrotoluenes (DNTs) are often found as co-contaminants where TNT is manufactured. To deal with TNT contamination, many physico-chemical processes have been proposed and have achieved varying degrees of success. Granular activated carbon was an effective way to remove TNT in water, but it is an expensive and less efficient option for treating soil-bound contamination (Spain, Hughes et al. 2000, Moteleb, Suoidan et al. 2001). Incineration has also been used for treating TNT contaminated soils, however, the high cost and the environmental burdens of incineration (e.g., production of toxic ash and greenhouse gas emissions) limits its use at large scale (USEPA 2005).

Bioremediation of TNT contaminated sites has been proposed as an environmentally-friendly and cost-effective alternative to physico-chemical treatment (Dı́az, Ferrández et al. 2001, Gerth, Hebner et al. 2003, Gerth and Hebner 2007). Many microorganisms are able to cleave aromatic rings through oxygenase-catalyzed upper biodegradation pathways and assimilate its carbon skeletons for growth using central metabolic intermediates, but oxygenolytic biodegradation pathways are limited to mono- and di-nitroaromatic compounds (e.g., MNTs and DNTs) (Dı́az, Ferrández et al. 2001). The presence of three electron-withdrawing nitro substituents inserted on the aromatic ring of TNT restricts similar oxidative cleavage mechanism for TNT (Rieger, Sinnwell et al. 1999). As opposed to electrophilic attacks by oxygenases, reductive biotransformation of TNT to 2-hydoxylamino-4,6-dinitrotoluene (2-OHADNT), 2-amino-4,6-dinitrotoluene (2-ADNT), 4-amino-2,6-dinitrotoluene (4-ADNT), 2,4-diamino-6-nitrotoluene (2,4-DANT), and 2,6-diamino-4-nitrotoluene (2,6-DANT) has been frequently reported (Boopathy and Kulpa 1992, Duque, Haidour et al. 1993, Funk, Roberts et al. 1993, Pasti-Grigsby, Lewis et al. 1996, Fiorella and Spain 1997, Widrig, Boopathy et al. 1997), with strict anaerobic conditions leading to the production of 2,4,6-triaminotoluene (TAT) (Hawari, Halasz et al. 1998). Consequently, a number of bioremediation strategies based on reductive biotransformation of TNT and TNT derivatives have been implemented, including soil slurry reactors (Funk, Roberts et al. 1993, Montgomery, Coffin et al. 2013), composting (Boopathy, Manning et al. 1998, Achtnich, Sieglen et al. 1999), and land farming (Boopathy and Kulpa 1992). While reductive biotransformation of TNT has been promising, it is not by far an ultimate solution, as the toxic derivatives may still accumulate in living organisms (Lachance, Renoux et al. 2004, Kuperman, Checkai et al. 2005) or eventually be released into water via weathering processes (Kuperman, Checkai et al. 2005, Taylor, Lever et al. 2009). TNT denitration, i.e., the release of nitrite ions and formation of TNT-denitrated metabolites via reductive nucleophilic attacks by hydride ions catalyzed by hydride transferases (van Dillewijn, Wittich et al. 2008), was proposed as a prerequisite for subsequent oxidative cleavage of the aromatic ring. Only a rare study reported an oxidative microbial degradation pathway of TNT (Tront and Hughes 2005), which generates the metabolic intermediate 3-methyl-4,6-dinitrocatechol after TNT denitration. In this study, ^14^C-labeled TNT degradation in combination with the production of ^14^C-labeled CO2, organic metabolites and nitrite was observed in laboratory microcosm incubations constructed with soil samples from a historically contaminated site located in Wisconsin (Tront and Hughes 2005, Akkaya, Nikel et al. 2019).

Although there is lack of significant evidence of an oxidative pathway for mineralization of TNT, a series of test plots were setup at the historically contaminated site in Wisconsin (Barksdale TNT plant) where periodic tilling was implemented, and the levels of different nitroaromatics were monitored. In the Barksdale TNT plant study, TNT contaminated soil plots were aerated through tilling from 2007 to 2012. Tillage was carried out each season, while it was performed more often (once in a month) during the summer from June to September. Chemical analyses of samples from this study showed significant decreases in the concentration of TNT (Figure 1) and DNTs (not shown) was observed in tilled soils. Both the historical contamination of nitroaromatics and the process of tillage likely had a significant impact on the soil microbiota composition and function. Prior studies have shown that anthropogenic activities, such as tilling, can change the physical and chemical properties of the soil thereby affecting the soil bacterial community structure (George, Liles et al. 2009, Kihara, Martius et al. 2012, Lupwayi, Lafond et al. 2012, Navarro-Noya, Gómez-Acata et al. 2013). Navarro-Noya et al. (2013) investigated the effect of tillage, soil physical and chemical properties, and crop-rotation on soil bacterial community structure. At the phylum level, relative abundances of *Actinobacteria*, *Beta-Proteobacteria* and *Gamma-Proteobacteria* were affected by the tillage treatment in this study. Soil total organic carbon (TOC), total nitrogen, and pH, affected the relative abundance of *Bacteroidetes*, *Beta-Proteobacteria*, *Cyanobacteria* and *Gemmatimonadetes*. In another work, the effect of TNT contamination on microbial community was compared with pristine soil (George, Eyers et al. 2008). Denaturing gradient gel electrophoresis (DGGE) of 16S rRNA gene amplicons was applied to study bacterial community composition in TNT contaminated soil and pristine soil samples. TNT contamination caused a shift in bacterial community composition, with *Pseudomonadaceae* and *Xanthomonadaceae* as the dominant families detected in TNT contaminated soil. While bacterial community structure in soil has been shown to change as a result of mechanical aeration and TNT contamination separately, no studies have investigated the effects of both.

**Figure 1.**
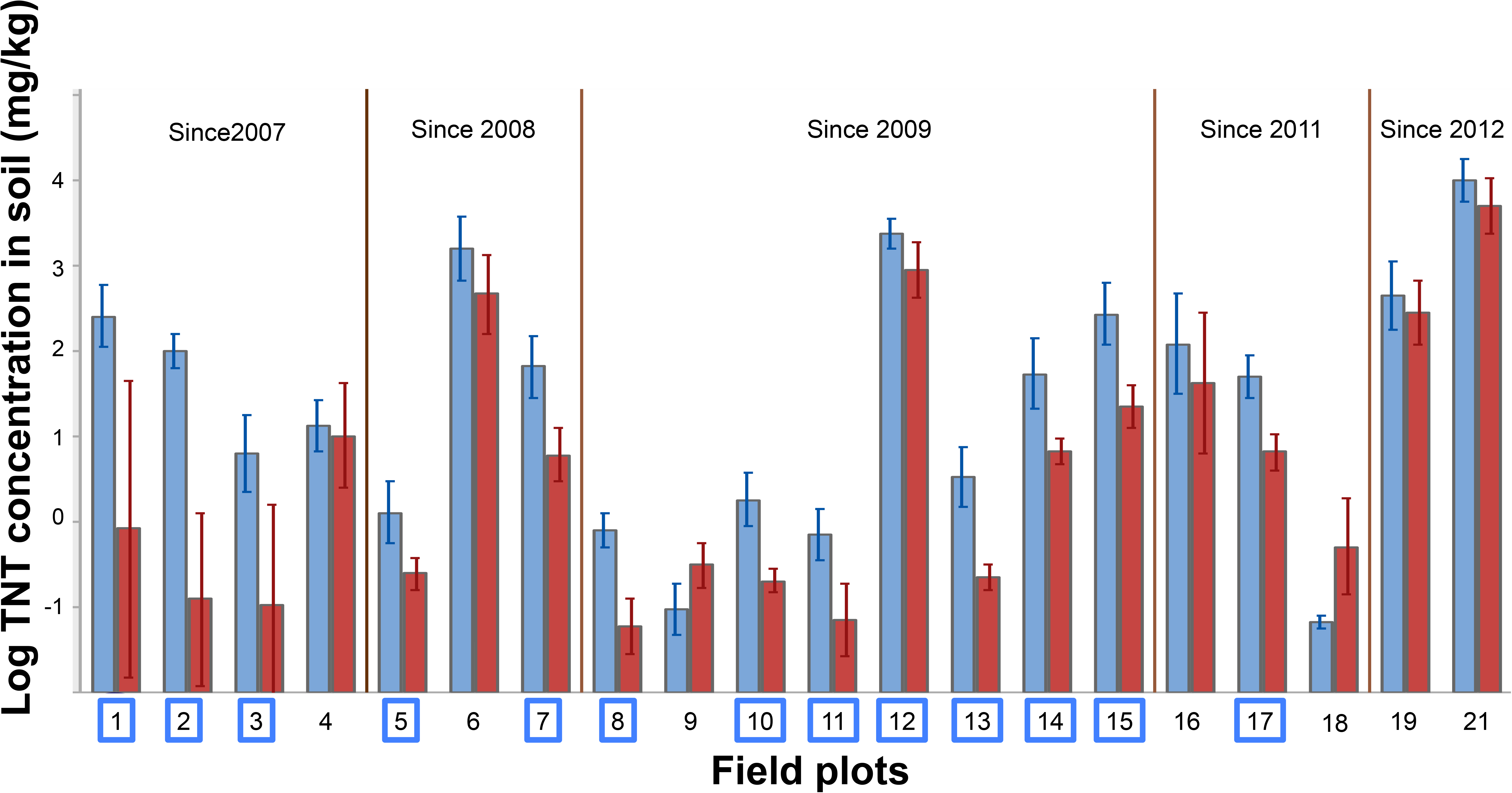
TNT concentration in soil for each field plot, in its first year (2007) of mechanical tillage (blue bars) vs. in 2013 (red bars, except for plot 9 whose last measurement was in 2012). Many field plots show significant TNT loss (plot numbers in blue boxes).

In this study, high-throughput sequencing of phylogenetic marker genes was applied to elucidate the microbial community structure of bacteria under the impact of historical nitroaromatics contamination and subsequent periodic tillage. Samples were collected from untilled-contaminated soils, tilled-contaminated soils, and pristine soils from the former Barksdale TNT manufacturing plant. In addition to the taxonomic study, the metagenomic composition of nitroaromatic-contaminated soil under tillage treatment was investigated to determine which metabolic pathways could play a role in degradation of nitroaromatics. A customized pipeline incorporating metagenomic binning approaches helps us recover genetic information of key populations of the microbial community that exist in tilled, nitroaromatics contaminated soils.

## 4. Materials and Methods

### 4.1. Field-scale Mechanical Tillage Experiments

Soil samples used for DNA extraction were collected in triplicates on a single date in 2012 from the former Barksdale TNT plant at three different field sites. Vegetation clearance was performed prior to the construction of the field plots. The first site represented a pristine area (“Pristine”) that was not contaminated by nitro-aromatic explosives. The second site had previously contained a ditch that received TNT-containing storm water from nearby buildings where TNT was manufactured (“Untilled + TNT”). The third site was a 8.5 m by 10.9 m field plot that received wastewater discharge from TNT manufacturing processes (“Tilled + TNT”), which was not tilled before 2008. In order to be able to compare the untilled and tilled samples with TNT contamination, the two sites chosen are located in close proximity to each other, with the same soil characteristics (structure and texture) and similar TNT concentrations. Construction of the field plots consisted of clearance of vegetation and debris (including large pieces of TNT and DNT crystals) followed by mechanical tillage to improve aeration of the soil. The tiller could reach an effective depth of 56 cm. The plots were managed to avoid saturation of the soils and subsequent anoxic conditions. Surface water was prevented from entering the plot by shallow soil mounds constructed of topsoil piled and compacted along the uphill sides of the plot. Storm water from heavy rains was managed using sedimentation traps and silt fencing. To keep storm water from saturating the pore space of the tilled material, drain tiles were installed below the tilled depth from the test plots to a trap where the water was contained and eventually evaporated. Typically, the tillage events were carried out four times a year, usually once per month from June to September, if weather permitted. In the first year of plot construction and tillage, samples for the quantification of nitroaromatic explosive compounds, nitrate, nitrite, pH and soil moisture were collected after each tilling event. After the first-year samples were collected at the beginning and end of the field season, usually June and September. Nitroaromatic explosives were quantified using the EPA 8330, EPA 8321A, and EPA 8270 methods. At the time of sample collection for the study (June 2012) the plot had been tilled for five years.

### 4.2. Sampling and DNA Extraction from Field Test Plots

Soil samples were collected from the top 15.2 cm of soil. Samples were shipped overnight on ice and stored at -20 °C until processing. The amount of soil used for DNA extraction ranged, from 4.5 to 10.5 g (no sieving of the soils was performed). DNA was isolated using the MoBio® PowerMax Soil DNA Isolation kit (Carlsbad, CA). The concentration of isolated DNA ranged from 2.3 to 30.3 ng/μL in a total elution volume of 5 mL, as measured by a Nanodrop 1000 spectrophotometer (Thermo Scientific, Wilmington, DE). Isolated DNA was stored at -20°C until further processing for amplicon sequencing.

### 4.3. Illumina HiSeq 2000 Sequencing of 16S rRNA Gene

Triplicate polymerase chain reaction (PCR) amplification was performed for each sample to minimize jackpot effects and PCR biases. PCR primers targeted the 16S rRNA gene V4 hypervariable region (see PCR specifications and PCR primer design in Supplementary Information). PCR products from each individual sample were combined and purified with 1.5% agarose gel electrophoresis. DNA was recovered using the Ultra-Clean® GelSpin® DNA Extraction Kit (Mo Bio Laboratories, Inc., Carlsbad, CA). For each individual sample, PCR products were quantified using Nanophotometer P-300 (Implem, Westlake Village, CA) and Nanodrop ND-3300 (Thermo Scientific, Inc., Waltham, MA). Individual PCR products with unique barcodes were mixed in equimolar ratios and the pooled samples, with the associated sequencing primers, were sent to the California Institute for Quantitative Biosciences (QB3 facility, Vincent J. Coates Genomics Sequencing Laboratory, University of California, Berkeley). Quality control using Qubit® 2.0 Fluorometer (Life Technologies, Grand Island, NY) and quantitative PCR was conducted by the QB3 facility. Illumina® HiSeq 2000 sequencing was performed for 150-bp nucleotide paired-end multiplex sequencing, according to the manufacturer’s instructions. A control lane with balanced base genomic libraries was set to increase nucleotide diversity.

### 4.4. Sequence Assembly

Paired-end Illumina reads were assembled using Fast Length Adjustment of SHort reads (FLASH, version1.2.6) (Magoč and Salzberg 2011). Overlap between paired-end reads of 150 bp was set to range between 5 bp to its full length, with a maximum mismatch density of 50%. The assembled sequences were then demultiplexed in the open source software package Quantitative Insights Into Microbial Ecology (QIIME, version 1.7.0) (Caporaso, Kuczynski et al. 2010), using maximum consecutive low-quality bases of 5, minimum consecutive high-quality bases of 60% of the original reads, maximum N’s of 5 and the default Phred quality, threshold of 3. Taxonomic assignment was performed in QIIME. For the 16S rRNA gene, the assembled sequences were assigned to Greengenes (version13_5) OTUs at a threshold of 97% pairwise identity (McDonald, Price et al. 2012).

### 4.5. Taxonomical Analysis

Alpha diversity of each sample was calculated using the Simpson index (Simpson 1949). Beta diversity was analyzed by using principal coordinate analysis (PCoA) with weighted UniFrac method guided by the Greengenes reference tree of 97% OTUs (Lozupone and Knight 2005). This was done for all OTUs identified, as well as for OTUs belonging to the order *Burkholderiales*. Phylogenetic tree for the family *Alcaligenaceae* was trimmed from the Greengenes reference tree. Statistical analysis of different soils was performed using two-tailed *t*-test with the assumption of unequal variance (Welch *t*-test), and *p*-values were corrected using the FDR correction (Storey 2002). Statistical significance was defined as tests with a *p*-value and *q*-value both lower than 0.05. In addition, term frequency-inverse document frequency (TF-iDF) feature selection was applied to identify OTUs characterizing the pristine and untilled soil samples as described by Lan *et al*. (Lan, Kriete et al. 2013).

### 4.6. Sequence Accession

The 16S rRNA gene, ITS2 and metagenomic sequences are available in NCBI BioProject no. PRJNA235951.

### 4.7. Metagenomic Analysis of “Tilled + TNT”: Assembly, Binning, Gene Annotation and Taxonomical classification

Illumina reads were binned to investigate genes involved in degradation of TNT in the soil samples, as well as taxonomic classification of microbial community samples. Metagenomic studies were performed only for “Tilled + TNT” soil samples. There were three metagenomic libraries from the “Tilled + TNT” samples. Initially, the metagenomic data was filtered to remove low quality end of the reads using Sickle tool (Joshi and Fass 2011). Then, IDBA denovo assembly tool was used to assemble the whole metagenomic data (Peng, Leung et al. 2010). IDBA output contigs was used as reference for binning. BinSanity (Graham, Heidelberg et al. 2017), MaxBin2 (Wu, Simmons et al. 2015), MetaBAT (Kang, Froula et al. 2015), COCACOLA (Lu, Chen et al. 2017) and CONCOCT (Alneberg, Bjarnason et al. 2014) were used for binning of the metagenomic libraries. To improve the quality of the bins, DASTools was used to refine the bins of all five binning tools outputs (Sieber, Probst et al. 2018). For quality assessment of the bins, CheckM (Parks, Imelfort et al. 2015) was used to evaluate DASTool bins. DASTools bins were submitted to RAST (Aziz, Bartels et al. 2008) and Kbase (Arkin, Cottingham et al. 2018) for annotation against SEED database (Overbeek, Begley et al. 2005). In addition to 16S rRNA taxonomic classification of bacterial community, taxonomic classification of the metagenome assembled genome (MAG) was also performed by using CheckM, PhyloPhlAn (Segata, Börnigen et al. 2013) and CAT/BAT (von Meijenfeldt, Arkhipova et al. 2019). DNT degradation pathway is also studied exclusively.

The sequence similarity of putative *dntD* genes in MAGs to extradiol ring cleavage dioxygenases in NCBI was determined by using the CLUSTALW software package (Thompson, Higgins et al. 1994) for multiple sequence alignment of translated amino acid sequences and generating a phylogenetic tree using FastTree package (Price, Dehal et al. 2010).

## 5. Results and Discussion

### 5.1. TNT removal in soil tillage experiment

Although manufacturing of explosives ceased over 60 years ago at the Barksdale plant site, the high concentrations of TNT found in the investigated soil samples suggest that natural weathering and attenuation of TNT was minor. A recent tillage experiment started in 2007 on field plots at Barksdale demonstrated that periodic mechanical tillage could promote significant removal of TNT (Figure 1). The decrease of TNT concentration varies from plot to plot (Figure 1). Decreases in the concentrations of 2,4-DNT and 2,6-DNT were also observed. For example, in 2008 the mean concentrations for 2,4-DNT and 2,6-DNT were 530 and 70.6 mg/kg, respectively (95% confidence intervals of 177-884 and 17.7-123.5 mg/kg, respectively) while in 2012 the mean concentrations had decreased to 25.3 and 9.8 mg/kg, respectively (95% confidence intervals of 7.7-43 and 2.9-16.7 mg/kg, respectively). Over this period, concentration of 2-A-4,6-DNT and 4-A-2,6-DNT did not change significantly. During year 2012, combined nitrite/nitrate levels in the “Tilled + TNT” plot ranged from 12 to 73 mg/kg while soil pH ranged from neutral to slightly acidic (as low as 6.1). Active nitrate/nitrite, production can result in mild acidic soil, even if there are many other factors that could have affected pH (Fortner, Zhang et al. 2003, Han, Mukherji et al. 2011).

### 5.2. Bacterial community structure dynamics by 16S rRNA and metagenomic analysis

To explore the effects of nitroaromatic explosives contamination and gain new insights into possible effects of tillage and aeration on the contaminated soil native microbial community, a comparison microbial diversity of pristine soil, TNT contaminated soil (Untilled + TNT), and tilled TNT contaminated soil (Tilled + TNT) from the Barksdale site was carried out. Both DNT and TNT concentrations in the pristine soil were below the detection limit. DNT and TNT concentrations for TNT-contaminated soil treatments were acquired in 2012, the year when DNA extraction was performed. In the “Untilled + TNT” soil, TNT concentration was as high as 38,500 mg/kg, whereas it was an average of 2,595 mg/kg in the “Tilled + TNT” soil. In contrast, 2,4-DNT and 2,6-DNT concentrations in the soils were less different (32 mg/kg and 6 mg/kg in the untilled soil and 25 mg/kg and 10 mg/kg in the tilled soil). The lower concentrations of DNT in soils compared to TNT could be contributed to the fact that the primary manufacturing product was TNT (while DNT were undesired product), the relatively higher solubility of TNT (~100 mg/L) compared to DNT (~400 mg/L), and the fact that DNT is relatively more easily biodegraded as compared to TNT. As presented in Table 1, the biodiversity (measured by the Simpson index) was clearly reduced in the contaminated soil samples (both untilled and tilled) in comparison to the pristine soil samples, as previously described (Eyers, George et al. 2004). Using weighted UniFrac for beta-diversity measurement, we generated the principal coordinate analysis (PCoA) plot from the OTU abundances (Figure 2), which demonstrates clear separations between the samples of different field plots and clustering among samples from the same field plot. TNT contamination decreases the overall microbial community diversity in TNT contaminated soils compared to pristine soils.

**Table 1.** Biodiversity for each soil sample measured by the Simpson index.

| *Soil Type* | *Sample Name* | *Simpson Index* |
| --- | --- | --- |
| *Pristine* | P1 | 0.9867 |
|  | P2 | 0.9851 |
|  | P3 | 0.99 |
| *Tilled + TNT* | T1 | 0.8868 |
|  | T2 | 0.8917 |
|  | T3 | 0.7494 |
| *Untilled + TNT* | U1 | 0.8954 |
|  | U2 | 0.849 |
|  | U3 | 0.8667 |

**Figure 2.**
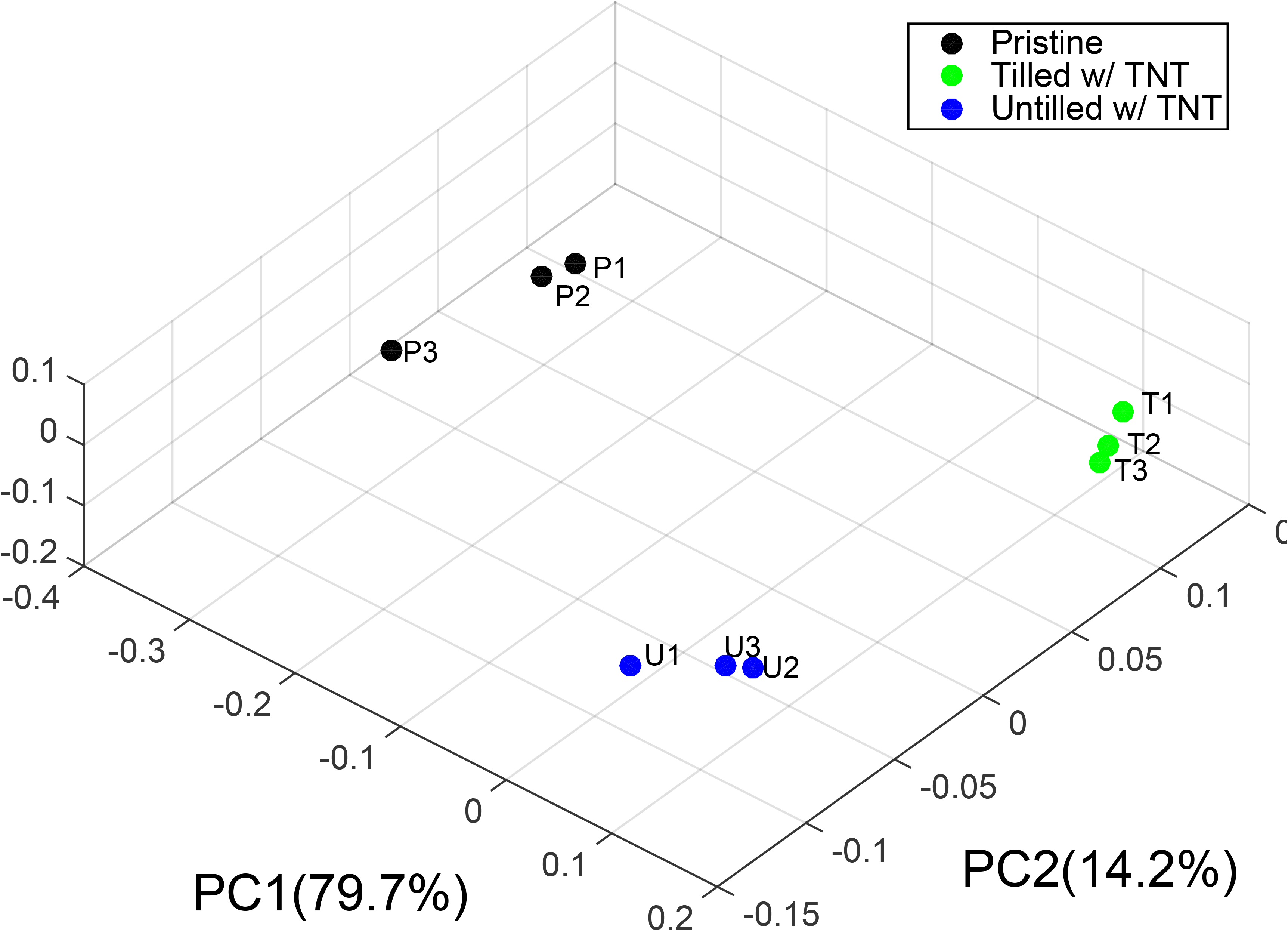
Principle coordinate analysis generated from the 97% OTU abundances, using weighted UniFrac.

In another work, the effects of TNT contamination on microbial communities was compared with pristine soil (George, Eyers et al.) using denaturing gradient gel electrophoresis (DGGE) of 16S rRNA gene amplicons. It was found that TNT contamination caused a shift in the bacterial communities. in this study, dominant bacterial populations in TNT contaminated soil were *Pseudomonadaceae* and *Xanthomonadaceae*.

The structural differences among the samples were further discriminated at different taxonomic levels (*i.e.*, phylum, family, genus, and species levels). Figure 3 shows the relative abundances of different phyla in the three soils. Although most of the 16S rRNA genes from the pristine soil were annotated as *Proteobacteria* (47.2 ± 13.2%) and *Acidobacteria* (25.9 ± 11.8%), the bacterial structure in this soil was more diverse than in the contaminated soils (Table 1), where more than 90% of the sequences belonged to *Proteobacteria*.

To further examine which microbial assemblages were dominant in the microbial communities of the tilled and untilled TNT-contaminated samples, we compared the top six most abundant families across the samples. *Burkholderiaceae* dominated the untilled soil (79.7 ± 3.0%) while *Alcaligenaceae* dominated the tilled soil (54.7 ± 9.8%), both belonging to the order *Burkholderiales* (Figure 4). Interestingly, a PCoA plot just using OTUs belonging to *Burkholderiales* (Figure 5) clearly separated the three soils into different clusters, indicating the structural changes of this order may be correlated with TNT contamination and tilling.

At the genus level, *t*-test analysis was again performed between the soil samples (Table S1). Several genera were found to be significantly increased in the tilled soil compared to the untilled or pristine soils, including *Achromobacter*, *KD1-23*, *Luteimonas*, *Microbacterium*, *Mycoplana*, *Pedobacter*, *Phaeospirillum*, *Pseudomonas*, *Salinibacterium*, *Thermomonas*, *Thiobacillus* and *Variovorax*. All these genera were relatively more abundant in the tilled samples, representing 31.5 ± 3.3% of the sequences detected the tilled soil combined while only 1.2 ± 0.4% in the pristine and 4.3 ± 2.1% in the untilled soils, indicating that they may be favored by aerobic conditions caused by mechanical tillage. Among them, *Achromobacter* from the *Alcaligenaceae* family was the dominant genus in the tilled samples (14.5 ± 3.4%). Consistent with the family level analysis, *Burkholderia* was the most abundant genus in the untilled, TNT-contaminated sample (78.7 ± 2.8%) and was significantly higher in the untilled soil compared to both tilled (0.1 ± 0.1%) and pristine (5.6 ± 1.9%) soils. Through term frequency-inverse document frequency (TF-iDF) feature selection, a technique recently shown to be effective to identify features relevant to particular sample traits in metagenomic data (Lan, Kriete et al. 2013), we examined the difference between the pristine and untilled TNT-contaminated samples at the genus level. We found that in addition to *Burkholderia*, there was a small presence of *Thauera* in the untilled TNT-contaminated soil, which has been shown to grow on toluene in aerobic and anaerobic conditions (Shinoda, Sakai et al. 2004).

**Figure 3.**
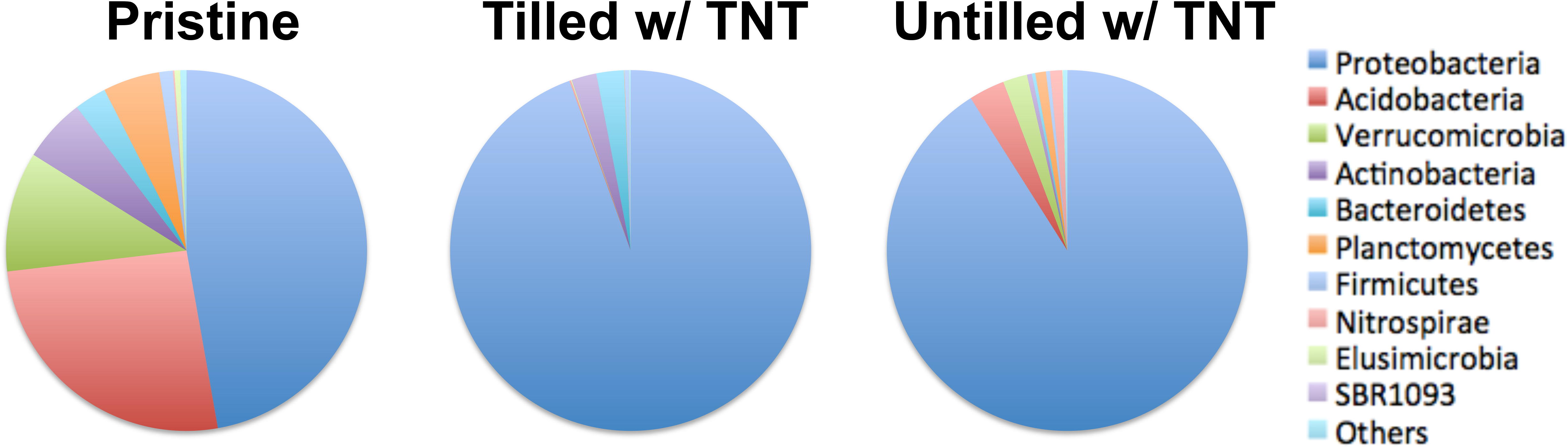
Relative abundances of different bacteria phyla in the three soils. While Proteobacteria dominates both the tilled and untilled soil, the remaining fraction of the microbial community is mostly composed of different phyla.

**Figure 4.**
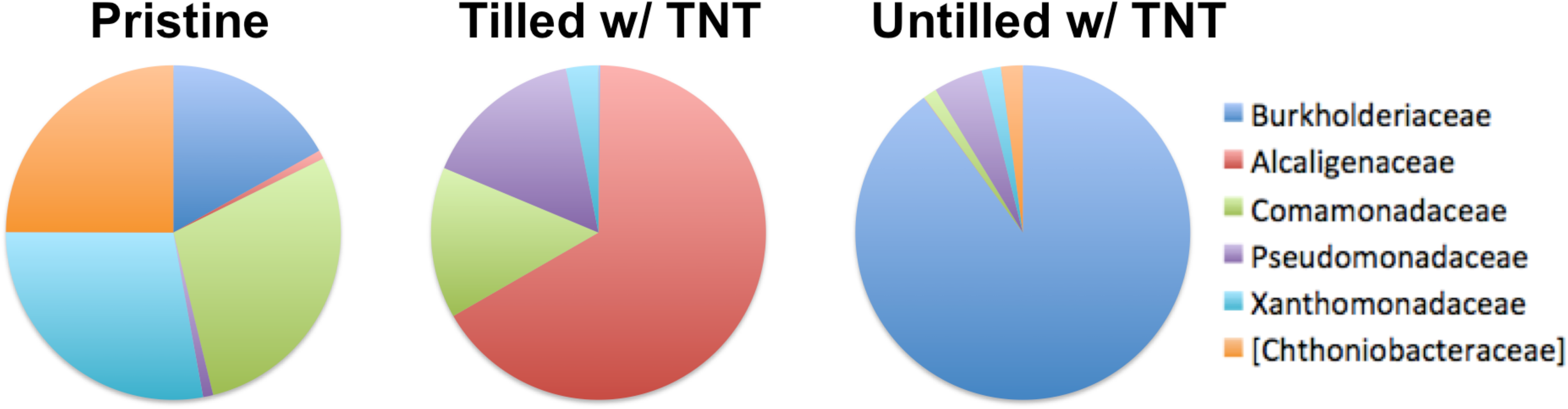
Relative abundances of the six most abundant bacteria families across different soils. Burkholderiaceae becomes the most dominant family that could survive TNT contamination, and after tilling, the relative abundance of Alcaligenaceae rapidly increased.

**Figure 5.**
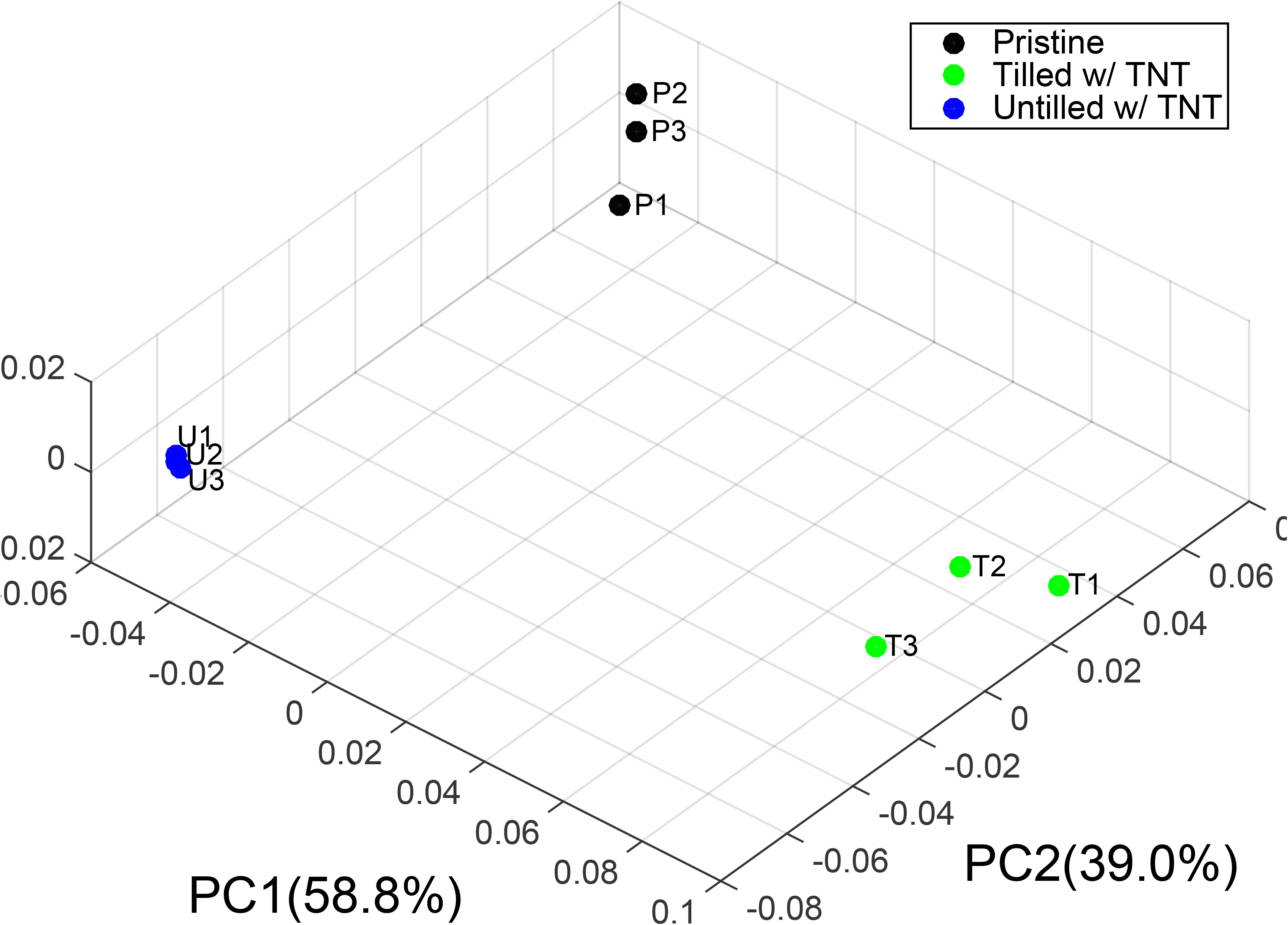
Principle coordinate analysis generated from the OTU abundances of Burkholderiales, using weighted UniFrac.

The most abundant OTU detected in the pristine soil was a *Xanthomonadaceae* (Greengenes OTU 105699) that accounted for only 7.0 ± 1.3% of the sample, demonstrating the high diversity observed in the pristine soil. In contrast, the two most abundant OTUs in the untilled TNT-contaminated soil belonged to the *Burkholderia* genus and accounted for 29.0 ± 6.0% (Greengenes OTU 668303) and 11.9 ± 1.7% (Greengenes OTU 125947) of the samples. The most abundant OTU in the tilled soil belonged to the *Alcaligenaceae* family (Greengenes OTU 565246), representing 35.3 ± 11.6% of the tilled sample sequences whereas this family only accounted for 0.1% of the untilled and around 0.2% of the pristine samples. A phylogenetic tree of *Alcaligenaceae* OTUs present (Figure 6) illustrates that this strain of interest is most closely related to *Achromobacter*.

**Figure 6.**
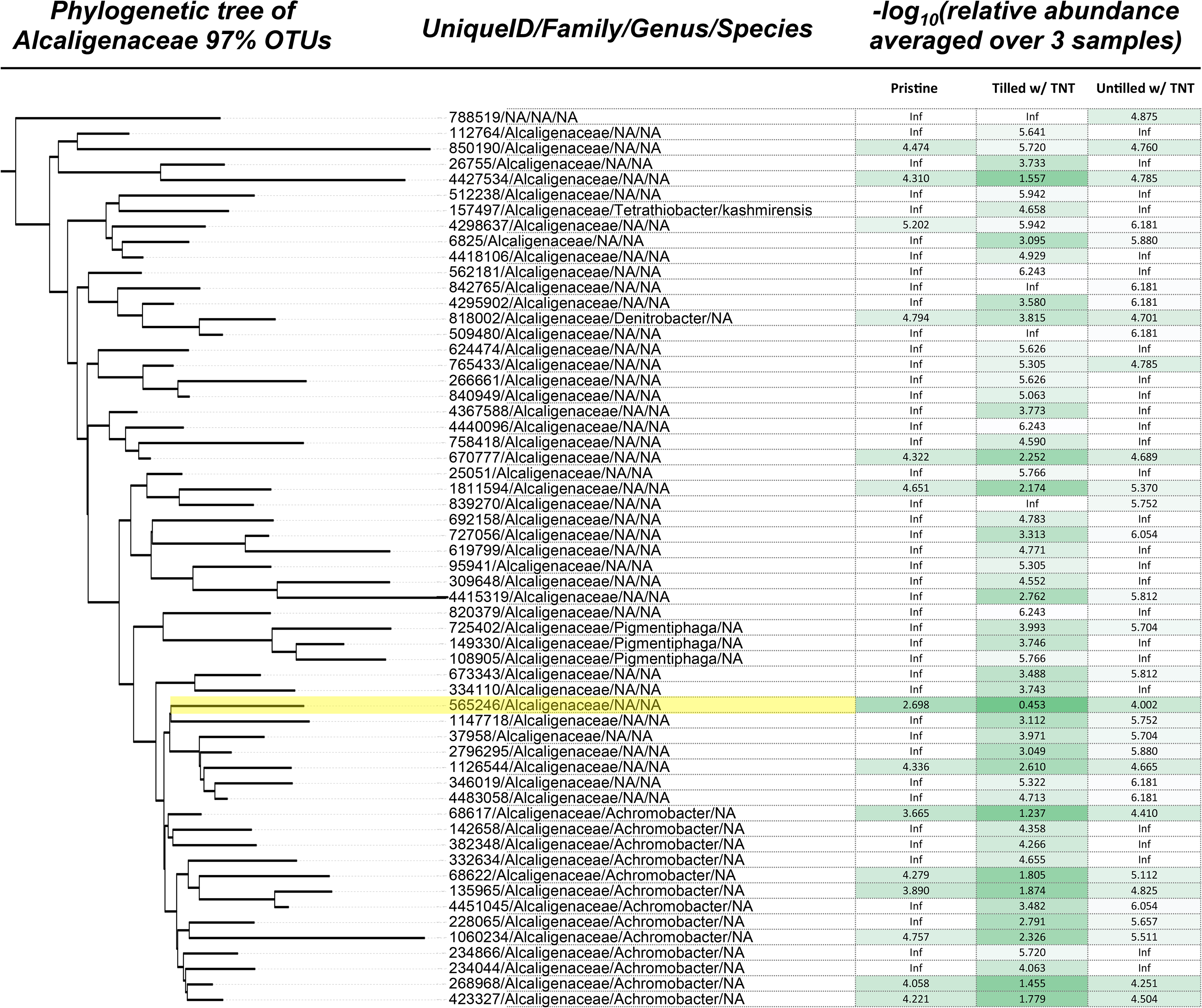
Phylogenetic tree and relative abundance of Alcaligenaceae OTUs identified in the soil samples. Negative log of relative abundance was shown for each OTU in the three soils, ranging from zero (green) to infinity (white). Leaf label for the most abundant OTU in the “Tilled + TNT” sample (Greengenes OTU 565246) was highlighted in yellow.

In addition to 16S rRNA analysis, metagenomic data used for taxonomical classification of “Tilled+TNT” soil plots. MAGs generated by DASTools and their related parameters are shown in Table S2. Twenty-two out of thirty-five bins had completeness more than 50%. Because, the contamination of generated bins was high, different approaches used to increase the reliability of the results. Phylogenetic classification of the MAGs was performed by CheckM, PhyloPhlAn and CAT/BAT. The results of MAGs phylogenetic classification are shown in Table S3. At the phylum level most of the metagenomic MAGs are classified as *Proteobacteria*. At genus level, a MAG is assigned to *Achromobacter* and another MAG is assigned to *Pseudomnas*, which confirms 16S rRNA based taxonomical classifications.

As observed in many bioremediation studies, the presence of contaminants is often associated with a decrease of biodiversity (species richness and evenness) and the presence of a few dominant phylotypes that can tolerate and/or take advantage of the selective pressure imposed by the contaminants (Smit, Leeflang et al. 1997, Ruberto, Vazquez et al. 2003, Vinas, Sabaté et al. 2005, Baek, Yoon et al. 2007). In our 16S rRNA gene analysis, we also found a correlation between the presence of TNT and a significant decrease in the overall biodiversity (Table 1 and Figure 2). In addition, *Proteobacteria* dominated the bacterial communities in both TNT-contaminated soils. This result is consistent with previous studies examining the effects of TNT contamination on soil microbial communities (Gong, Gasparrini et al. 2000, Siciliano, Gong et al. 2000, George, Eyers et al. 2008, Travis, Bruce et al. 2008, Limane, Muter et al. 2011).

Although *Proteobacteria* dominated both TNT-contaminated soils in this study, the microorganisms that were able to survive in the presence of TNT (*i.e.*, OTUs with high relative abundances) differed between untilled and tilled soils. In particular, the dominance of the *Burkholderiaceae* family in the untilled TNT-contaminated soil is remarkable. As the biodiversity decreased relative to the pristine samples, *Burkholderiaceae* became the most dominant microbial family in the untilled, TNT-contaminated soil, suggesting that it has high tolerance for TNT or is highly competitive in the presence of TNT. In the untilled, TNT-contaminated soils, about 99% of the *Burkholderiaceae* are from the genus *Burkholderia*, which is consistent with previous findings that some *Burkholderia* strains are chemo-attracted to TNT (Leungsakul, Keenan et al. 2005).

In contrast with the untilled soils that were dominated by *Burkholderia*, the most abundant OTU family in the tilled soils belonged to *Alcaligenaceae* (Greengenes OTU 565246). This strain, which is closest to the genus *Achromobacter*, was highly abundant in the tilled soil, while only composing a fairly small amount of the pristine and untilled soils. In addition, *Achromobacter* OTUs in total accounted for 14.6 ± 3.5% of the tilled soils, while less than 0.1% in the other two soils. *Achromobacter* sp. DNT (formerly *Burkholderia* sp. DNT) and *Achromobacter* sp. NDT3 has been reported to mineralize 2,4-DNT and *Achromobacter xylosoxidans* degrades 2,6-DNT (Parales, Spain et al. 2005, Hudcova, Halecky et al. 2011, Perez-Pantoja, Nikel et al. 2013). BLAST the 16S rRNA gene from the *Alcaligenaceae* strain we identified, against the non-redundant NCBI database revealed a 98% percent identity match to the 16S rRNA gene in *Achromobacter xylosoxidans AU1011* (GI: 15384334), suggesting that this particular genus was involved in the removal of TNT. In a study by Gumuscu and Tekinay (2013), a novel strain of *Achromobacter spanius* STE11 identified which was capable of producing DNT and using TNT as the only nitrogen source over a wide range of pH (4.0-8.0). It is possible that the increased presence of *Achromobacte*r *sp.* observed in the “Tilled + TNT” soil reflects a role in the observed decreases in concentrations of TNT and DNT isomers.

### 5.3. Metagenomic data analysis, genes and annotation

While taxonomic analyses indicated which OTUs are dominant in tilled, TNT-contaminated soils, further analysis of the metagenomic data could reveal what functional capabilities (i.e., gene content) of certain microorganisms exist under tilled, TNT-contaminated conditions. A, customized pipeline to cluster reads from the metagenomic libraries into MAGs was used to functionally characterize the gene content of “binned” populations in the microbial community, as well as taxonomically classify each MAG. Then, each individual MAGs were examined for the presence of specific genes associated with nitroaromatic transformation, aromatic ring metabolism, and nitrogen metabolism. The major focus of the functional analysis of the bins was on previously proposed biotransformation pathways of DNT and TNT (Hughes, Wang et al. 1998, Esteve-Núñez, Caballero et al. 2001, Johnson, Jain et al. 2002), which are discussed separately in the sections below.

#### 5.3.1. TNT transformation

Although MNT and DNT compounds can be oxidatively cleaved, the symmetrical arrangement of nitro groups on TNT makes the aromatic ring electron-deficient and electrophilic, which in turn prevents oxidative attacks of TNT using electrophilic-driven oxidation by oxygenases. This arrangement favors reductive, nucleophilic transformation mechanisms (Stenuit, Eyers et al. 2009), under both aerobic and anaerobic conditions. There are three major scenarios proposed for initial TNT transformation: (i) reduction of nitro moieties, (ii) production of Meisenheimer complexes of TNT through nucleophilic attacks by hydride ions, and (iii) Bamberger Rearrangement. Following these initial transformation scenarios for TNT, the products of the initial step can undergo different intermediate steps before the aromatic ring is cleaved and completely mineralized. These Intermediate Steps and Mineralization pathways are described below. In our work, metabolic pathways of the aforementioned scenarios are used as reference to construct possible TNT transformation pathways present in the microbial community of the tilled, TNT-contaminated soil based on the gene content of our MAGs.

##### 5.3.1.1. Initial Transformation Scenarios

###### 5.3.1.1.1. Reduction of nitro moieties of TNT

Reduction of nitro moieties of TNT could proceed through both aerobic and anaerobic conditions (Esteve-Núñez, Caballero et al. 2001). A few studies (Crawford 1995, Ederer, Lewis et al. 1997) showed that the nitro (-NO2) moieties of TNT can be reduced to nitroso (-NO), hydroxylamino (-NHOH) and eventually amino (-NH2) groups. Since this nitroreduction could happen to any of the three nitro moieties on TNT, many, different combinations of reduced products can form. Complete reduction of all nitro groups in TNT eventually results in triaminotoluene (TAT), only under strict anaerobic conditions. When nitro groups are completely reduced to amino groups, they are believed to dissociate from the aromatic rings (Boopathy and Kulpa 1994, Vorbeck, Lenke et al. 1994, Crawford 1995, Ederer, Lewis et al. 1997, Esteve-Núñez, Caballero et al. 2001).

Genes contents of MAGs were investigated to find out possible scenarios in TNT degradation (Figure 7 and Table 2). Genes encoding for nitroreductases that catalyze reduction reactions on nitroaromatic compounds were detected in our MAGs (Gao, Ellis et al. 2009). There are two different types of the nitroreductases: oxygen-sensitive (type II) and oxygen-insensitive (type I). Some experimental studies (Mason and Holtzman 1975, Peterson, Mason et al. 1979) showed that TNT are reduced by both types of nitroreductases and also by hydride transferase type I and type II (van Dillewijn, Wittich et al. 2008, Stenuit and Agathos 2010). Oxygen-sensitive nitroreductases (type II) reduce the nitro groups of TNT only under anaerobic condition (Somerville, Nishino et al. 1995, Esteve-Núñez, Caballero et al. 2001, Roldán, Pérez-Reinado et al. 2008). Oxygen insensitive nitroreductases (type I), on the other hand, could reduce nitro groups of TNT under both aerobic and anaerobic condition. Sequence alignment of nitroreductases genes in our MAGs showed that only type I NAD(P)H nitroreductase enzymes are present. The presence of genes for type I nitroreductases means that reduction of nitro groups on TNT could proceed throughout the cycling between aerobic and anaerobic condition due to periodic tilling.

**Table 2.**
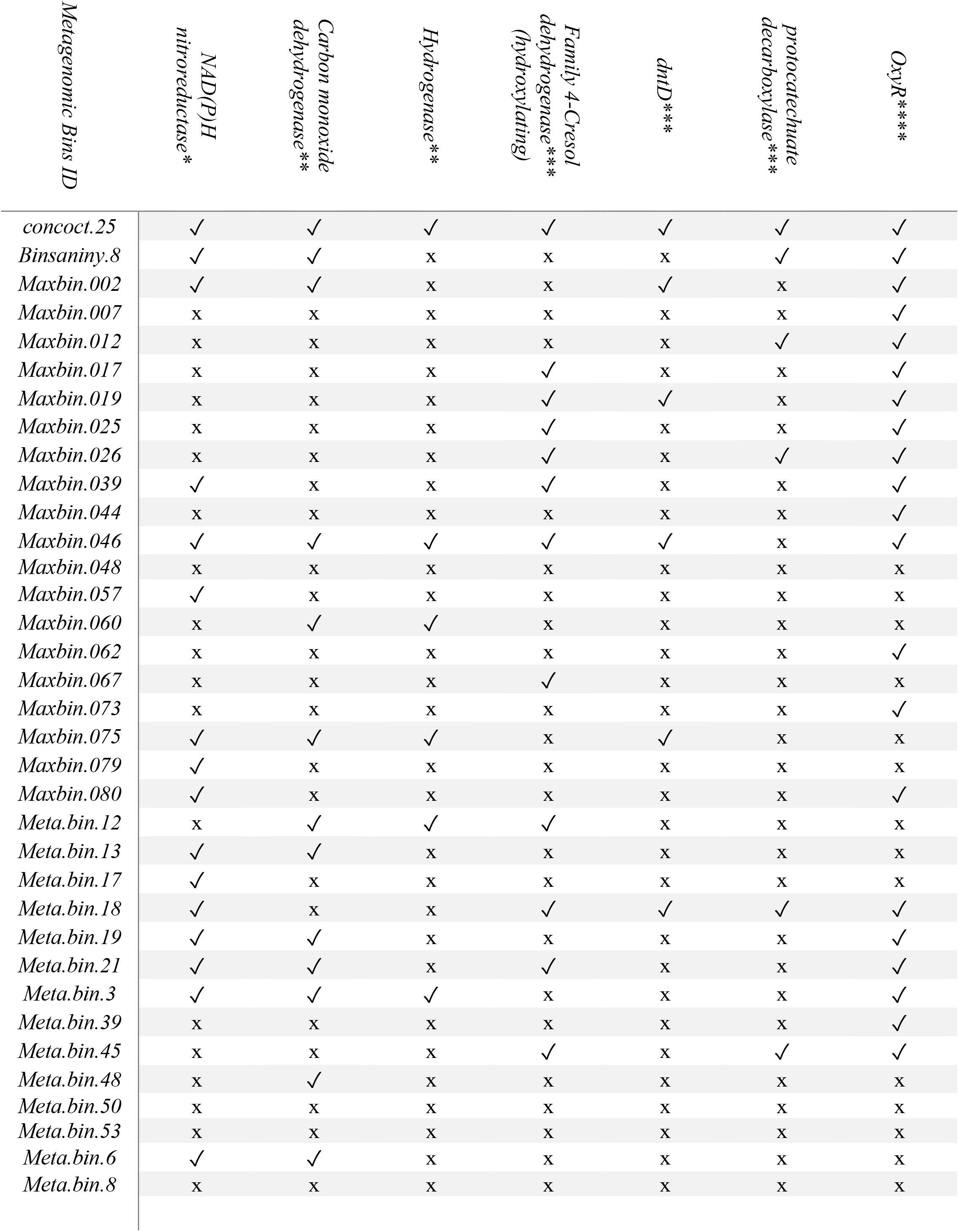
Key genes in TNT and DNT transformation in MAGs for tilled, TNT contaminated soil. * Initial Transformation genes; ** initial Bamberger Rearrangement genes; *** Intermediate Transformation; **** Intermediate Transformation

**Figure 7.**
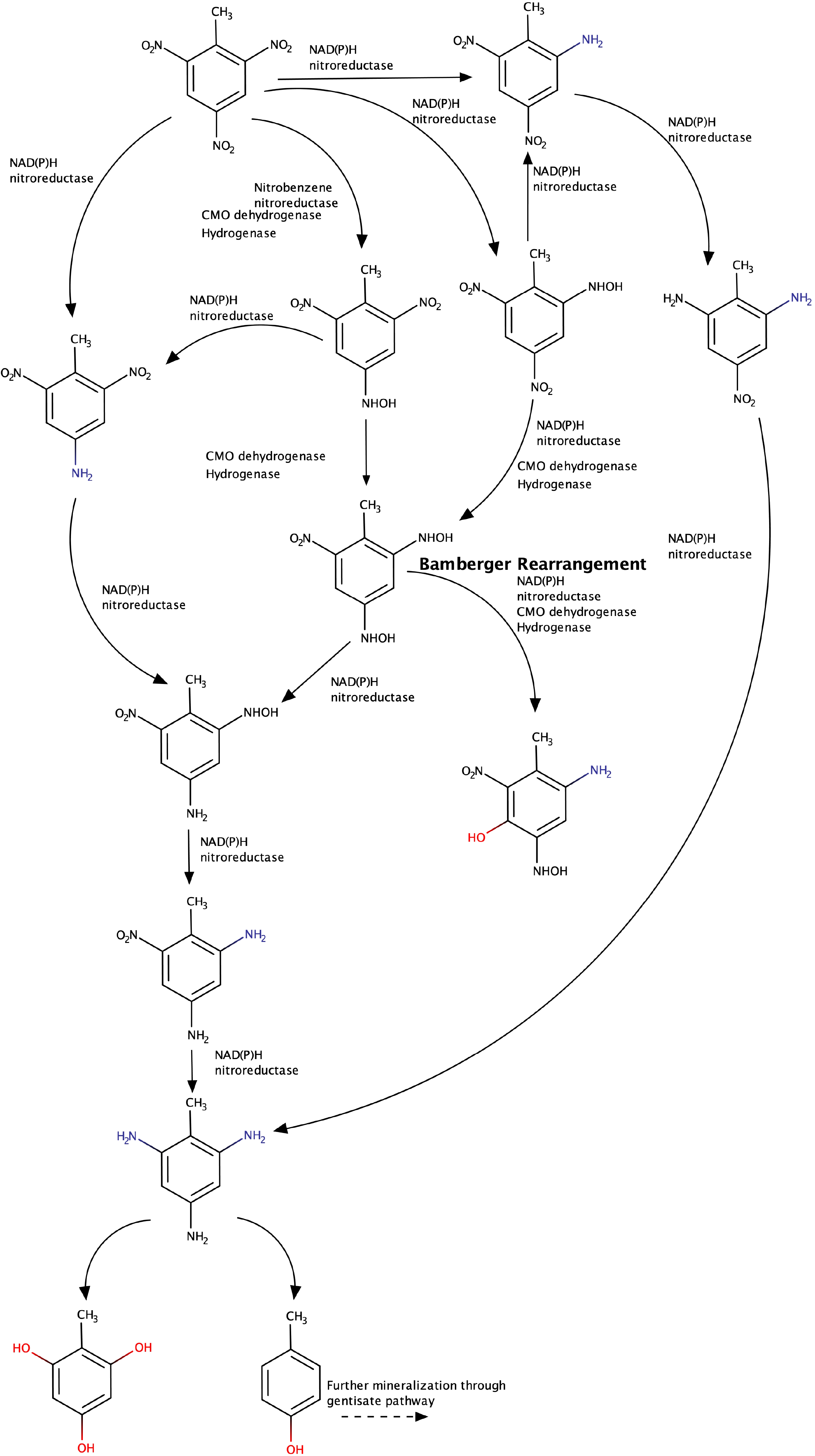
Initial transformation pathway of TNT based on gene content of constructed MAGs in tilled, TNT contaminated soil.

###### 5.3.1.1.2. Meisenheimer complexes

The electron-withdrawing inductive (-I) and resonance (-M) effects of the nitro group makes the aromatic ring of TNT prone to nucleophilic attacks to generate reaction adducts, such as hydride Meisenheimer complexes (McFarlan 1999, Esteve-Núñez, Caballero et al. 2001, Gao, Ellis et al. 2009). Formation of monohydride and dihydride Meisenheimer complexes of TNT (H^—^TNT and 2H--TNT) under aerobic conditions have been confirmed experimentally (Vorbeck, Lenke et al. 1994, French, Nicklin et al. 1998) and can lead to the removal of a nitro group leading TNT-denitrated metabolites. Under aerobic conditions, the formed dinitrotoluene could transform to nitrotoluenes, which can be, further metabolized (Gao, Ellis et al. 2009). The nitro groups of the dinitrotoluenes formed from the Meisenheimer complexes could undergo nitroreduction, as described above or degrade aerobically as described below. We investigated two enzymes have been reported to play a key role in transformation of TNT to TNT-denitrated metabolites through Meisenheimer complexes. One of these key enzyme which catalyzes formation of Meisenheimer nitroreductases is pentaerythritol tetranitrate (PETN) reductase (French, Nicklin et al. 1998). Another enzyme which have been investigated in this study is NADH-dependant flavoprotein oxidoreductase (XenB). This enzyme reduces TNT by formation of dihydride Meisenheimer complexes (Blehert, Fox et al. 1999). In-depth analysis of our constructed MAGs showed that there are no genes attributed to PETN reductase or XenB. This means that Meisenheimer complexes formation could not be a possible scenario in initial transformation of TNT in the Tilled+TNT contaminated soil plots in this study.

###### 5.3.1.1.3. Bamberger Rearrangement

The Bamberger Rearrangement pathway is an alternative TNT transformation pathway that is presumed to be only possible under strictly anaerobic conditions (Hughes, Wang et al. 1998, Ahmad and Hughes 2000). In this pathway, after the reduction of nitro groups and formation of dihydroxylaminonitrotoluene (DHANT) under either aerobic or anaerobic conditions, DHANT could transform under strict anaerobic conditions to 2-amino-4-hydroxylamino-5-hydroxyl-6-nitrotoluene and hydroxylamine-4-amino-5-hydroxyl-6-nitrotoluene (Figure 7). In the Bamberger Rearrangement pathway, TNT is reduced by hydrogenase and carbon monoxide dehydrogenase enzymes by way of ferredoxin or methyl viologen (Huang, Lindahl et al. 2000). We investigated the presence of the genes encoding for these Bamberger Rearrangement genes in our constructed MAGs (Table 2). Although key genes (carbon monoxide dehydrogenase, monooxygenase and hydrogenase) in this pathway were found in a number of our MAGs (Table 2), they were not obligate anaerobic species such as *Desulfovibrio* spp., *Clostridium pasteurianum* or *Clostridium thermoaceticum* (Boopathy, Kulpa et al. 1993, Khan, Bhadra et al. 1997, Hughes, Wang et al. 1998, Huang, Lindahl et al. 2000). The genera classification of the MAGs containing these, Bamberger Rearrangement genes are *Phenylobacterium, Methylibium, Acidovorax, Pseudomonas, Erythrobacter, Parvibaculum, Nitrospira, Caldimonas, Oceanibaculum* and *Nocardioides*. We searched for any publicly available genomes in NCBI belonging to genera that possess these key Bamberger Rearrangement genes (carbon monoxide dehydrogenase, monooxygenase and hydrogenase). We BLASTed the key genes from the MAGs against the NCBI genome data base. The results are summarized in (Table S4). The results indicate that at least one of the genes encoding these enzymes, is present in every genome of these genera (except *Caldimonas*) in NCBI data base (Table S4). Future experimental studies to validate the functionality of these genes that potentially encode the Bamberger Rearrangement enzymes from the aforementioned genera would be helpful for better understanding of nitroaromatic compounds transformations.

##### 5.3.1.2. Intermediate Transformation

After initial reduction of nitro moieties intermediate products of TAT or aminophenols are generated, respectively. Further transformation of these molecules up to aromatic ring cleavage, which we describe as “intermediate transformation steps” will be discussed here.

TAT, the intermediate products produced via nitroreductases, is an unstable molecule that has two different fates under aerobic and anaerobic conditions. Under aerobic conditions, TAT molecules can polymerize with other TAT molecules to form azo derivatives, (e.g., 2,2’,4,4’-TA-6,6’-azoT and 2,2’,6,6’-TA-4,4’-azoT). TAT azo derivatives are recalcitrant compounds that are considered as dead-end products in TNT degradation studies (Esteve-Núñez, Caballero et al. 2001). While under strictly anerobic and slightly acidic (pH 5-7) conditions, it has been reported that TAT transforms abiotically to 2,4,6 hyroxytoluene (THT) (Funk, Roberts et al. 1993). In our study, the pH of tilled, TNT contaminated soil is slightly acidic (pH 6.1) and there were periodic aeration and anaerobic conditions, which suggest that both TAT azo derivatives and THT could have been formed. It is hypothesized that THT could undergo ring cleavage and eventually be mineralized under aerobic conditions (Alneberg, Bjarnason et al. 2014, Sieber, Probst et al. 2018). Figure 8 is a pathway based on those proposed by Serrano et al (Serrano-González, Chandra et al. 2018) for, transformation of THT. In our study, the presence of genes associated with the aromatic ring cleavage pathway of THT in our constructed MAGs were found (Table 2 and Figure 8).

**Figure 8.**
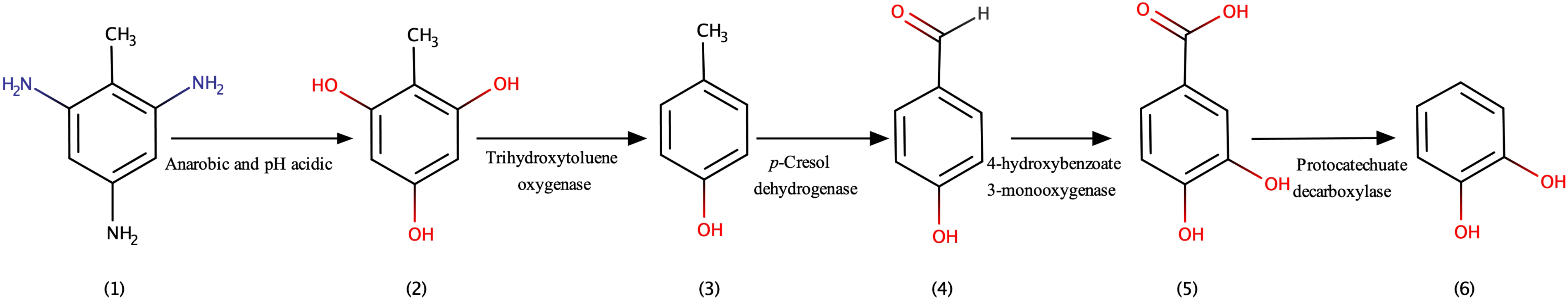
TAT degradation pathway based on the gene content of MAGs. TAT (1) transformatiom to 4-hydroxytoluene (2) by Trihydroxytoluene oxygenase enzyme. Then, methyl group on p-cresol (4-hydroxytoluene) oxidize and results 4-hydroxybenzoate. 4-hydroxybenzoic acid could oxidized to Protocatechuic acid. The next step in this pathway could be catechol (dihydroxybenzene).

The first step in this pathway is transformation of THT to 4-hydroxytoluene, which is catalyzed by a trihydroxytoluene oxygenase (Figure 8). This is a key reaction in initiating ring cleavage of THT (Haigler, Johnson et al. 1999). The gene encoding trihydroxytoluene oxygenase is called *dntD* and was only previously reported to be found in a *Pseudomonas* capable of degrading nitroaromatic compounds (Monti, Smania et al. 2005). We found that *dntD* exists in a number of our MAGs (concoct.25, Maxbin.002.sub, Maxbin.019.sub, Maxbin.046.sub, Maxbin.075.sub, Meta.bin.18), including those taxonomically classified as *Achromobacter* and *Pseudomonas* (Tables 2 and 3).

**Table 3.** The dnt genes group in DNT transformation in MAGs for tilled, TNT contaminated soil.

| *Metagenomic Bins ID* | *dntAa* | *dntAb* | *dntAc* | *dntAd* | *dntB* | *dntC* | *dntD* |
| --- | --- | --- | --- | --- | --- | --- | --- |
| *concoct.25* | ✓ | x | ✓ | x | ✓ | ✓ | ✓ |
| *Binsaniny.8* | ✓ | ✓ | ✓ | x | x | ✓ | x |
| *Maxbin.002* | ✓ | ✓ | x | x | ✓ | x | ✓ |
| *Maxbin.007* | x | x | x | x | x | x | x |
| *Maxbin.012* | x | x | x | x | x | ✓ | x |
| *Maxbin.017* | x | x | x | ✓ | x | x | x |
| *Maxbin.019* | x | x | x | ✓ | ✓ | ✓ | ✓ |
| *Maxbin.025* | x | x | ✓ | ✓ | x | x | x |
| *Maxbin.026* | x | x | x | ✓ | x | ✓ | x |
| *Maxbin.039* | x | x | x | ✓ | x | x | x |
| *Maxbin.044* | x | x | x | x | x | x | x |
| *Maxbin.046* | x | ✓ | ✓ | ✓ | ✓ | x | ✓ |
| *Maxbin.048* | x | x | x | x | x | x | x |
| *Maxbin.057* | x | x | x | x | x | x | x |
| *Maxbin.060* | x | ✓ | ✓ | x | x | x | x |
| *Maxbin.062* | x | x | x | x | x | x | x |
| *Maxbin.067* | x | x | x | ✓ | x | x | x |
| *Maxbin.073* | x | x | x | x | x | x | x |
| *Maxbin.075* | ✓ | ✓ | ✓ | ✓ | ✓ | ✓ | ✓ |
| *Maxbin.079* | ✓ | x | x | x | x | x | x |
| *Maxbin.080* | ✓ | x | x | x | x | x | x |
| *Meta.bin.12* | x | ✓ | ✓ | ✓ | x | x | x |
| *Meta.bin.13* | ✓ | ✓ | x | x | x | x | x |
| *Meta.bin.17* | ✓ | x | x | x | x | x | x |
| *Meta.bin.18* | ✓ | x | x | ✓ | ✓ | ✓ | ✓ |
| *Meta.bin.19* | ✓ | ✓ | x | x | x | x | x |
| *Meta.bin.21* | ✓ | ✓ | x | ✓ | x | x | x |
| *Meta.bin.3* | ✓ | ✓ | ✓ | x | x | x | x |
| *Meta.bin.39* | x | x | x | x | x | x | x |
| *Meta.bin.45* | x | x | x | ✓ | x | ✓ | x |
| *Meta.bin.48* | x | ✓ | x | x | x | x | x |
| *Meta.bin.50* | x | x | x | x | x | x | x |
| *Meta.bin.53* | x | x | x | x | x | x | x |
| *Meta.bin.6* | ✓ | ✓ | x | x | x | x | x |
| *Meta.bin.8* | x | x | x | x | x | x | x |

The Trihydroxytoluene oxygenase enzyme converts trihydroxytoluene to *p*-cresol. Themethyl group on *p*-cresol (4-hydroxytoluene) can then be oxidized by *p*-cresol dehydrogenase resulting in 4-hydroxybenzoate as the final product. The gene encoding for the *p*-cresol dehydogenas enzyme is also detected in our MAGs (Table 2). The produced 4-hydroxybenzoate can be converted to 4-hydroxybenzoic acid, which could be enzymatically oxidized to protocatechuic acid (dihydroxybenzoic acid) by protocatechuate decarboxylase. Protocatechuate decarboxylase oxidizes protocatechuic acid to catechol (dihydroxybenzene). A number of the enzymes involved in converting THT to catechol exist in the MAGs (Table 2 and TableS5). In addition to our proposed pathway, there many are alternative pathways for ring cleavage of aromatic compounds that hypothetically could lead to complete mineralization of THT. For example, the ring cleavage of hydroxybenzoate could occur through the Gentisate pathway. The gentisate 1,2-dioxygenase gene (*gtdA*) was found in Maxbin.046 (*Pseudomonas*) and Maxbin.075 (*Achromobacter*). The *gtdA* gene was also found in publicly available genomes of *Pseudomonas aeruginosa* and *Achromobacter xylosoxidans*. Other alternative pathways that we searched for in our MAGs for ring cleavage of THT intermediates are those listed in SEED Subsystems as “Metabolism of Aromatic Compounds” are shown in Table S5.

#### 5.3.2. DNT degradation

Transformation of DNT molecules are also another subject of our study because of its presence in the contaminated soils at the site. It was shown that DNT concentration decreased overtime in tilled soil and could potentially be due to biotransformation of DNT by microorganisms. Biological reductive transformation of DNT has been experimentally shown to proceed via nitroreductases similar to TNT described above (Kalafut, Wales et al. 1998). In addition, the asymmetrical structure of DNT makes direct oxidation under aerobic conditions—and hence aromatic ring cleavage—easier. Direct oxidation of DNT is catalyzed by a series of oxidative pathway enzymes and regulators encoded by *dnt* genes (Johnson, Jain et al. 2002, de las Heras, Chavarría et al. 2011, Perez-Pantoja, Nikel et al. 2013, Akkaya, Pérez-Pantoja et al. 2018). This aerobic transformation pathway catalyzed by dioxygenases involves the oxidation and removal of the two nitro groups, released as nitrite ions, eventually leads to ring cleavage (Figure 9). DNT oxidation is started by a multi compound hydroxylation dioxygenase (*dntA*). Then, methylnitrocatechol is oxidized by a monooxygenase (*dntB*) to a methylquinone. This compound reduces to trihydroxytolune by 2-hydroxy-5-methylquinone reductase (*dntC*), and eventually oxidize by THT oxygenase (*dntD)*. Our MAGs were investigated for the aforementioned *dnt* genes and the results are summarized in Table 3. The *dnt* genes were observed in MAGs, which are associated to *Pseudomonas, Phenylobacterium* and *Achromobacter*. Previous studies also indicate that some strains of *Burkhorderia,* such as *Burkholderia* sp. strain DNT (Haigler, Johnson et al. 1999) and *Achromobacter,* such as *Achromobacter* sp. NDT3 (Hudcova, Halecky et al. 2011) are able to metabolize DNT.

**Figure 9.**
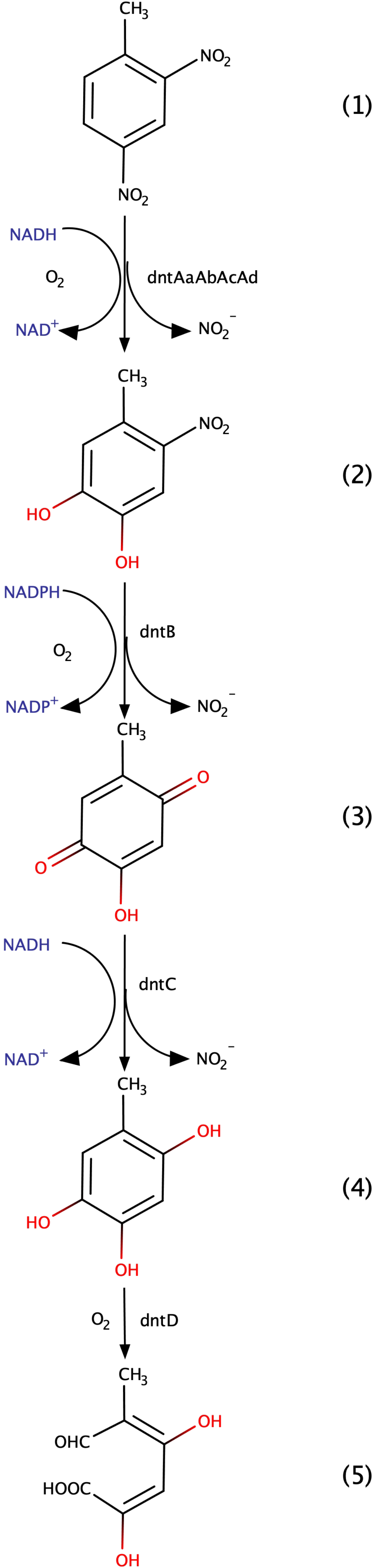
DNT degradation pathway based on the gene content of MAGs. DNT (1) oxidation is started by a multi compound hydroxylation dioxygenase (dntA). methylnitrocatechol (2) is oxidized by a monooxygenase (dntB) to a methylquinone (3). This compound reduces to trihydroxytolune (4) by 2-hydroxy-5-methylquinone reductase (dntC), and eventually oxidize by THT oxygenase (dntD) to 2,4-dihydroxy-5-methyl-6-oxo-2,4-hexadienoic acid (5).

Further interrogation of the putative *dntD* genes found in the MAGs was carried out to ensure they were more similar to 2,3,5-trihydroxytoluene (THT) 1,2-dioxygenase gene (*dntD*) found in *Burkholderia* sp. Strain DNT than other extradiol cleavage gene family I enzymes (Haigler, Johnson et al. 1999). The species name associated with accession number AF076848 provided for the *dntD* gene referenced in Haigler et al. (Haigler, Johnson et al. 1999) was *Burkholderia cepacia* (not strain associated with DNT). Amino acid sequences of the *dntD* genes detected in the MAGs were aligned with extradiol ring cleavage enzyme sequences from *Burkholderia cepacia* and other closely related species. A phylogenetic tree was created to determine the proximity of our MAGs putative *dntD* genes to other extradiol ring cleavage enzymes. Figure 10 indicates that the putative *dntD* sequences from Metabin.18, Maxbin.46 and Maxbin.75 were more closely related to the *dntD* enzyme in *Burkholderia cepacia*, supporting their likely involvement in the observed DNT degradation in the tilled samples.

**Figure 10.**
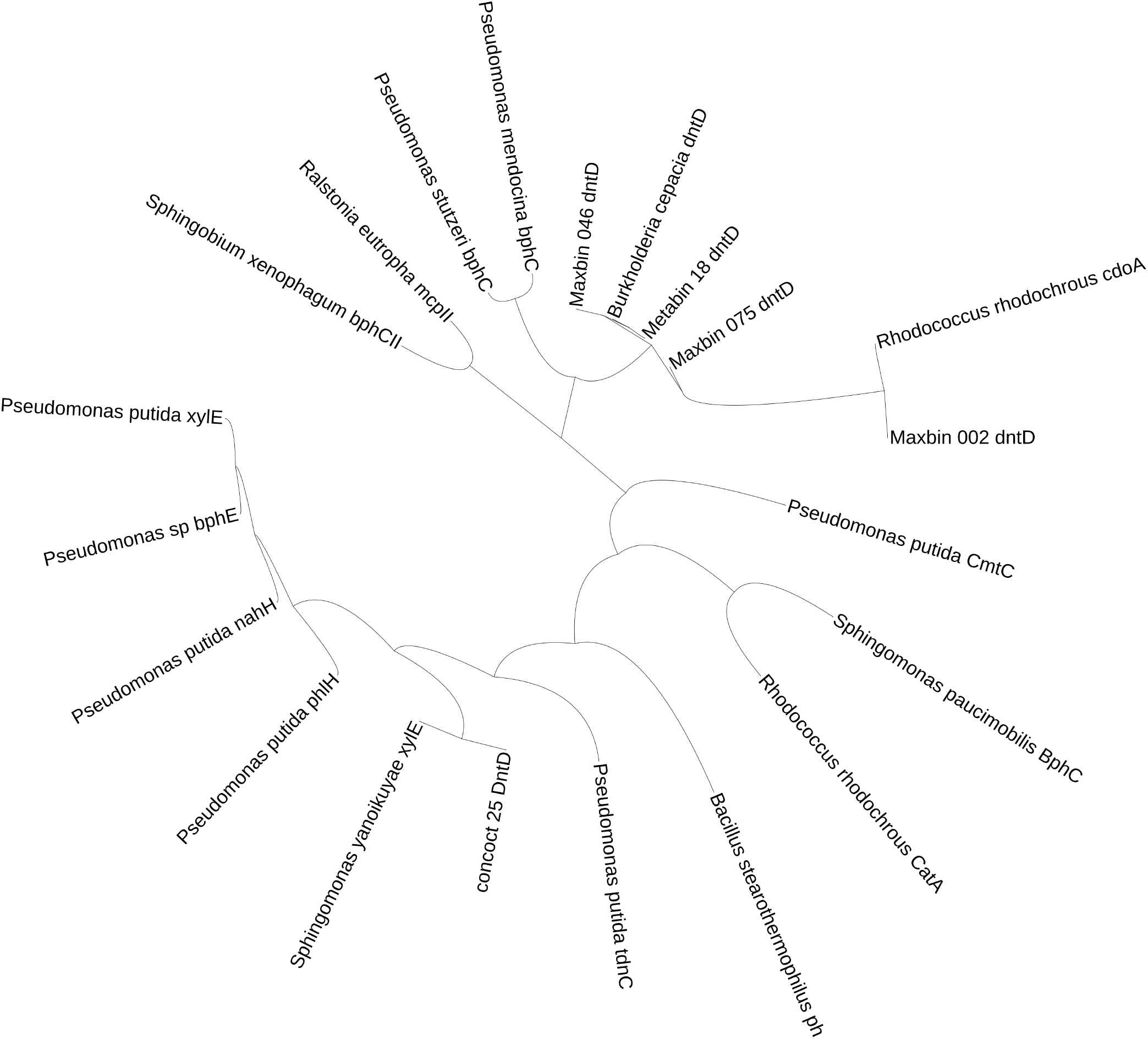
Phylogenetic tree showed proximity of MAGs putative dntD genes to other extradiol ring cleavage enzymes.

#### 4.3.3. Importance of OxyR and nitrogen assimilation genes in tilled, nitroaromatic contaminated soils

In addition to the genes that are directly involved in TNT and DNT transformations, we also examined microbial functions related to nitrogen assimilation of the ammonium and nitrite ions that are cleaved from TNT and DNT, as well those related to the ability of microbial populations to adapt to periods of aerobic condition in the tilled, TNT contaminated soils.

One of the genes investigated in this study is *OxyR*, a hydrogen peroxide-inducible activator. One by-product of the aerobic reactions discussed earlier is hydrogen peroxide that are toxic for living organisms. Microorganisms adopted a defense mechanism to overcome hydrogen peroxide (Lee, Godon et al. 1999, Chiang and Schellhorn 2012, Akkaya, Pérez-Pantoja et al. 2018). These microorganisms have groups of genes that reduce hydrogen peroxide to water. These genes are activated with *OxyR* which is a regulatory gene. *OxyR is* activated with high concentrations of hydrogen peroxide and starts translation of hydrogen peroxide reduction genes. In-depth analysis show that *OxyR* genes are present in number of our MAGs (Table 2). Presence of *OxyR* genes in our constructed MAGs is an indication of oxidative transformation activities in the soil. This is in agreement with our previous results that proposed pathways of complete aerobic degradation of DNT molecules and partial aerobic degradation of TNT molecules.

In addition to *OxyR*, we also examined nitrogen assimilation genes. Based on the proposed pathways for TNT and DNT transformation, nitrogen could be released in the form of nitrite groups or ammonia groups. TNT transformation is proposed to proceed through reduction of nitro groups in either aerobic or anaerobic conditions; While, nitro groups are released from DNT molecules under aerobic conditions. Investigation of MAGs shows presence of ammonia assimilatory genes as well as nitrate or nitrite reducing genes in a, number of the MAGs (Table S6). MAGs attributed to *Achromobacter* (Maxbin.75) and *Pseudomonas* (Maxbin.46) genera have genes encoding “Nitrate and nitrite ammonification” and “Ammonia assimilation” enzymes in SEED data base. Gumuscu and Tekinay (2013) identified a strain of *Achromobacter spanius STE11* capable of TNT and DNT transformation as the only nitrogen source over a wide range of pH (4.0-8.0). However, the *Achromobacter* strain only has an 83% 16S rRNA sequence identity to the strain we identified via our 16s rRNA amplicon libraries. It is possible that the increased presence of *Achromobacter sp.* observed in the “Tilled + TNT” soil reflects a role in the observed decreased concentrations of both DNT isomers and TNT.

## 6. Conclusions

In this study, TNT and DNT removal has been observed in field-scale experiments following periodic tilling of historically contaminated soils. Concomitantly, the microbial community structures of uncontaminated pristine soils, untilled contaminated soils, and tilled contaminated soils were investigated using high-throughput sequencing platforms. In addition, shotgun metagenome libraries of samples from tilled contaminated soils were generated. The major results gleaned from the microbial community data indicate a significant shift of the bacterial community at the family level between tilled and untilled contaminated soils, with tilled soils being dominated by *Alcaligenaceae* and untilled soils by *Burkholderiaceae*. At the genus level, *Acidovorax, Pseudomonas* and *Achromobacter* are dominant genera in “Tilled +TNT” soil (Table S1), with some members of the latter two genera reported as having the ability to degrade nitroaromatic compounds (Boopathy, Manning et al. 1994, Boopathy, Wilson et al. 1994, Somerville, Nishino et al. 1995, Parales, Spain et al. 2005, Hudcova, Halecky et al. 2011, Perez-Pantoja, Nikel et al. 2013).

In-depth metagenomic analysis of samples from tilled contaminated soils indicate the presence of genes that encode for enzymes that potentially could lead to mineralization of TNT and DNT under mixed aerobic and anaerobic periods. Determination of MAGs from the metagenomic data allowed us to examine which microbial populations could potentially be involved in the different steps of TNT and DNT mineralization pathways. In addition, the presence of *OxyR* regulatory genes that protect cells from oxygen radicals in MAGs that can catalyze the oxidative reactions within the intermediate transformation steps of TNT and aerobic DNT transformation. MAGs were also, identified that could assimilate nitrate and ammonia from the aforementioned transformation of TNT and DNT in the tilled contaminated soils. Confirmation that TNT or DNT can be used by the microbial communities as a nitrogen source could be investigated by isotope labeling of carbon and nitrogen in TNT and DNT.

## Acknowledgements

We thank E.E. Mack formerly with DuPont Corporate Remediation Group, C. Pooler and S. Larson from AECOM, and B. Nave from Chemours Corporate Remediation Group for their contribution to site monitoring and sample collection.

The work performed by the Drexel Ecological and Evolutionary Signal-processing and Informatics (EESI) lab (Y. Lan and G.L. Rosen) was also supported in part by a National Science Foundation (NSF) CAREER award number #0845827, NSF award number #1120622, and Department of Energy (DOE) Office of Science (BER) award DE-SC0004335.

We thank J. Tremblay and S.G. Tringe of the Department of Energy Joint Genome Institute (DOE JGI, Walnut Creek, CA) for the design and optimization of 16S rRNA gene and ITS2 primers for targeted Illumina sequencing. This work used the Vincent J. Coates Genomics Sequencing Laboratory at UC Berkeley, supported by NIH S10 Instrumentation Grants S10RR029668 and S10RR027303.

